# Myo-inositol in the Dorsal Anterior Cingulate Cortex is Associated with Anxiety-to-Eat in Anorexia Nervosa

**DOI:** 10.1101/2024.05.29.596476

**Authors:** Yulu Song, Sarah H. Guo, Christopher W. Davies-Jenkins, Angela Guarda, Richard A.E. Edden, Kimberly R. Smith

## Abstract

Anorexia nervosa (AN) is a mental and behavioral health condition characterized by an intense fear of body weight or fat gain, restriction of food intake resulting in low body weight, and distorted body image. Substantial research has focused on general anxiety in AN, but less is known about eating-related anxiety and its underlying neurobiological mechanisms. We sought to characterize anxiety-to-eat in AN and to examine neurometabolite levels in the dorsal anterior cingulate cortex (dACC), a brain region putatively involved in modulating anxiety-related responses, using edited magnetic resonance spectroscopy. Sixteen women hospitalized with AN and 16 women of healthy weight without a lifetime history of an eating disorder (healthy controls; HC) completed a computer-based behavioral task assessing anxiety-to-eat in response to images of higher (HED) and lower energy density (LED) foods. The AN group reported greater anxiety to eat HED and LED foods relative to the HC group. Both groups reported greater anxiety to eat HED foods relative to LED foods. The neurometabolite myo-inositol (myo-I) was lower in the dACC in AN relative to HC. In the AN group only, myo-I levels negatively predicted anxiety to eat HED but not LED foods and was independent of body mass index, duration of illness, and general anxiety. These findings provide new insight into the clinically challenging feature of eating-related anxiety in AN, and indicate potential for myo-I levels in the dACC to serve as a novel biomarker of illness severity or therapeutic target in individuals vulnerable to AN.

## Introduction

Anorexia nervosa (AN) is a mental and behavioral health condition characterized by intense fear of body weight or fat gain, severe restriction of food intake resulting in low body weight, and distorted self-perception of body shape or weight ^1^. AN has the second highest mortality rate of all psychiatric illnesses ^2^ and affects individuals of all ages, sexes, and genders ^3,4^. Biological females are particularly vulnerable, with age of onset occurring around puberty ^5^. While there are no pharmacological treatments for AN, weight restoration to a Body Mass Index (BMI) of 19-21 kg/m^2^ remains the mainstay of treatment for severe AN. Even so, relapse rates in the first year following weight restoration range from 30-50% ^6,7^. The mechanisms that maintain the disorder and underlie these high relapse rates remain unknown.

Anxiety is a risk factor shown to exacerbate disease severity and promote relapse in AN ^8^. Meal-associated anxiety is a striking clinical feature in patients with AN, especially when confronted with pressure to consume higher energy density (HED) foods^9^. Consequently, individuals with AN consume largely, if not exclusively, low calorie foods and limit consumption of calorically dense foods ^10–16^. While substantial research has focused on general anxiety in AN, less is known about eating-related anxiety and its underlying neural mechanisms.

The characterization of neural mechanisms of anxiety in mammals has largely emerged from circuit-based approaches in animal models^17,18^. Electrophysiological, lesion, and infusion studies have revealed that conditioned fear expression involves the amygdala, the ventral medial prefrontal cortex (vmPFC) including the infralimbic and prelimbic subregions, and the hippocampus^19–21^. In humans, structural and functional magnetic resonance imaging (MRI) studies have implicated the amygdala, insula, bed nucleus of the stria terminalis (BNST), and the prefrontal cortex (PFC) including the ventral PFC (vPFC) and the anterior cingulate cortex (ACC) in the development and expression of fear and anxiety responses. ^22–29^. Among these regions, the dorsal anterior cingulate (dACC), the human homolog of the rodent prelimbic cortex, is a relatively large cortical region that can be assayed by magnetic resonance spectroscopy (MRS) and is implicated in both negative threat appraisal and modulation of fear and anxiety responses and in functional MRI studies of anorexia nervosa.

Electrodermal skin conductance, a measure of sympathetic nervous system activity, has been shown to positively correlate with ACC blood flow^30^, dACC activation to a conditioned fear stimulus^31^, and dACC cortical thickness^31^ in healthy individuals, whereas lesions to the ACC are associated with impaired electrodermal activity in response to negative valence physical and psychological stimuli^32^. Functional MRI studies indicate that individuals with generalized anxiety disorder show greater ACC activation ^26,27^ and increased functional connectivity among the amygdala, vPFC, and ACC ^28,29^. Individuals with AN relative to controls show increased activation to high calorie food images in the dACC and ACC broadly ^33,34^, reductions in dACC gray matter volume^35^ and decreased blood perfusion of the ACC ^36–38^ although these appear to reverse, at least partially, with weight restoration ^39^. Taken together, these studies highlight involvement of the dACC in fear and anxiety-related conditions and in anorexia nervosa. MRI techniques are unable however to offer insights into the neurobiological mechanisms underlying these clinically challenging features of AN.

Proton MRS is a widely-used, noninvasive neuroimaging technique for examining levels of neurometabolites in the human brain *in vivo*. It quantifies concentrations of various metabolites within a region of interest (ROI) by detecting the magnetic properties of protons unique to each compound’s molecular structure ^40,41^. Although limited in number, previous MRS studies investigating metabolite levels in the dACC of individuals with AN relative to healthy controls have yielded mixed findings. These inconsistencies may be attributable to methodological differences across studies including magnet strength, inclusion of underweight versus weight-restored AN condition, and assessments in adolescents versus adults^42–47^. Additionally, measurement of low-concentration brain metabolites (e.g., gamma-aminobutyric acid (GABA), glutathione (GSH)) with conventional MRS is challenging due to limited separation and significant overlap of the metabolite peaks^40^. Recent methodological advances in our laboratory however have enabled edited MRS to simultaneously measure levels of multiple low-concentration, coupled metabolites. Briefly, the Hadamard Editing Resolves Chemicals Using Linear-combination Estimation of Spectra (HERCULES)^48^ editing scheme enables the extraction of distinct edited spectra from different Hadamard combinations. No study to date has investigated metabolite levels in individuals with AN using HERCULES.

We used a novel behavioral computer-based paradigm to measure anxiety-to-eat and the HERCULES editing scheme with MRS to quantify dACC neurometabolite levels in women with AN and in healthy weight women without a lifetime history of an eating disorder (healthy controls; HC). We hypothesized that women with AN would show greater anxiety to eat HED foods relative to lower energy density (LED) foods and greater anxiety-to-eat in general relative to HC. Consistent with previous MRS data acquisition in the ACC and dACC of adults with AN, we hypothesized that the AN group would show lower levels of myo-inositol (myo-I)^47^ and glutamate and glutamine (Glx)^43,47^, but similar levels of N-acetyl aspartate (NAA)^43,45,47^ total choline (tCho)^43,45^, and total creatine (tCr)^43,45,47^ relative to levels observed in healthy controls. Given the superiority of HERCULES editing scheme to measure low-concentration metabolites and its first application in the context of AN, we also compared GABA+ and GSH levels between groups as well as N-acetyl aspartyl glutamate (NAAG) and lactate (Lac). Exploratory analyses were then conducted to determine relationships between anxiety-to-eat and levels of neurometabolites in the dACC that significantly differed between groups.

## Materials and Methods

### Participants

Participants were recruited from an ongoing prospective observational study investigating anxiety-to-eat in individuals with AN and in HC (K01MH127178). Data collection occurred between September 22, 2022 and November 30, 2023. The protocol was approved by the institutional review board at the Johns Hopkins School of Medicine (IRB00214417). Women with AN were recruited from the Johns Hopkins Hospital during the initial phase of inpatient treatment, and HCs were recruited from the local community. For women with AN, eligibility criteria included the following: met DSM-V criteria for AN; currently hospitalized in the Johns Hopkins Inpatient Eating Disorders Program; 13-70 years of age; able to read, speak, and comprehend English fluently; no MRI contraindications including pregnancy, claustrophobia, metal implants, and history of brain injury. For HCs, eligibility criteria included: a BMI between 19-24.9 kg/m^2^; 13-70 years of age; no lifetime history of an eating disorder or axis I disorder; able to read, speak, and comprehend English fluently; no MRI contraindications as listed above. All participants provided written informed consent prior to study assessments.

### Measures

#### Overview

Participants completed two study visits (average interval of 5.03 days ± 0.79 days between visits) and were instructed not to consume any food or beverages apart from water after midnight the day prior to each visit. A glucose fingerstick was conducted to confirm that participants arrived fasted (<100 mg/dL). At both visits, participants reported interoceptive and appetite ratings on a visual analogue scale (VAS) prior to data collection. At Visit 1, participants completed a computer-based assessment of anxiety-to-eat on a 13-inch Macbook Air computer. Body weight and height were collected using a Tanita MC-980U plus body composition scale for individuals in the HC group and gowned morning weight was extracted from electronic medical records for individuals in the AN group. A link to complete psychological inventories was emailed to participants following completion of the first study visit. At Visit 2, all participants completed the scanning protocol which included a 10-minute MRS scan of the dACC. Scan day gowned body weight and height were extracted from electronic medical records for individuals in the AN group. Assessments are described in detail below.

#### Interoception Assessment (Visit 1 and 2)

Participants reported their levels of hunger, thirst, fullness, desire to eat, anxiety, and stress on a 145 mm visual analog scale (VAS), where “0” (left scale anchor) represented the least degree ever experienced and “+100” (right scale anchor) represented the greatest degree ever experienced of the respective variable of interest.

#### Anxiety-to-Eat Assessment (Visit 1)

Participants completed a computer-based behavioral task featuring 10 higher energy-dense (HED; 1.9–4.9 kcal/gram) food images and 10 lower energy-dense (LED; 0.2–1.4 kcal/gram) foods images. Each food was presented at a standard serving size (e.g., 1 banana, 1 glazed donut) and displayed on a 10-inch white plate. For each food image, participants were given the following instructions, “For each question, imagine you haven’t eaten in 4 hours. Rate the level of anxiety you would feel to eat the portion size of the food presented in the picture on a scale from 0 to +100, where 0 represents "no anxiety / least anxiety ever experienced" and +100 represents "most anxiety ever experienced".

#### Neuroimaging Protocol (Visit 2)

Data were collected on a Philips 3T Ingenia Elition RX scanner (Philips Healthcare, Best, the Netherlands) equipped with a 32-channel receive-only head coil. The scanning protocol included a 1 mm^3^ isotropic T1 MPRAGE (TR/TE=6.5/2.9ms, Flip angle=8°; 211 slices; voxel size = 1 mm^3^ (isotropic voxels) for MRS voxel placement and segmentation. The MRS voxel size was 30 (AP) x 30 (CC) x 30 (RL) mm^3^ in dACC, as shown in Figure 1. MR spectra were acquired using the following parameters: PRESS localization; TR/TE=2000/80 ms; 256 transients; HERCULES editing with 20-ms editing pulses applied at: 4.58, 4.18, and 1.9 ppm; 2048 datapoints; 2 kHz spectral width; VAPOR water suppression; and interleaved water reference correction ^48^. The dACC voxel was individually placed medially with the bottom of the voxel aligned with the top margin of the body of the corpus callosum (Figure 1).

**Figure 1.**
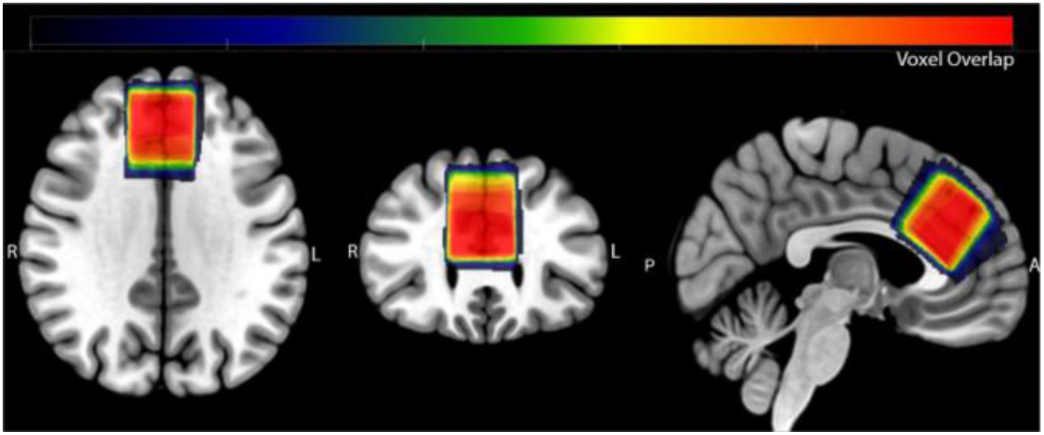
Location of the dACC voxels. The native space binary voxel mask for each participant was normalized to MNI space and overlaid onto the spm152 template to display anatomical overlap in voxel positioning between participants. Warmer (red) colors denote areas of greater overlap across participants.

#### Psychological Inventories

Participants completed the State-Trait Anxiety Inventory (STAI)^49^ and the Eating Disorder Examination Questionnaire (EDEQ)^50^. The STAI is a self-report measure of the presence and severity of generalized state (STAI-S) and trait (STAI-T) anxiety. The EDEQ is a self-report measure that assesses the range, frequency, and severity of eating disorder behaviors and is comprised of four subscales (eating restraint, eating concern, shape concern, weight concern). Both the STAI and EDEQ have good psychometric properties ^51–54^ (e.g., validity and reliability).

### Data Analysis

Three HCs had BMI values outside the defined lean range of 19-24.9 (i.e., 18, 25, and 25) and were subsequently excluded from the dataset. From the remaining 27 HCs, 16 were selected to match the age distribution of the test group; birth month and year were matched as close as possible between groups. This approach was taken to control for observed age-related effects on neurometabolite levels as measured by MRS ^55,56^ and to equalize the sample size between groups. Notably, analyses including the full sample of 27 HCs were conducted and yielded similar results to the age-matched groups discussed below. One participant in each group did not complete the psychometric instruments (STAI, EDEQ).

Each participant provided an anxiety-to-eat score (0-100) for each food in the computer-based task. Scores were averaged within an individual across the 10 LED foods and across the 10 HED foods to generate a mean LED and HED score for each participant. Responses on the STAI and EDEQ were calculated as instructed in the original manuscripts. IBM SPSS Statistics version 28 and GraphPad Prism version 10 were used to analyze and plot the demographic, psychometric, and psychophysical data. An alpha level of 0.05 was predetermined for all analyses. A priori analyses of variance or chi-square tests were conducted to examine differences in demographics, STAI and EDEQ scores, anxiety-to-eat ratings, and neurometabolite levels in the dACC between groups. Multiple linear regression was conducted to obtain predictive models for the main variable of anxiety-to-eat for both HED and LED foods using a stepwise procedure to automatically select the significant predictors among the independent variables of myo-I levels (found to be significantly different between groups), BMI, general anxiety as assessed via the VAS anxiety scale on the scan day, and blood glucose level on scan day. Duration of AN illness was also entered into the model in the AN group only.

MRS data were analyzed using the open-source analysis toolbox Osprey (version 2.4.0) ^57^ within MATLAB R2022a. The analysis procedures followed consensus-recommended processing guidelines for MRS data ^58,59^. Briefly, standardized analysis steps in Osprey included eddy-current correction based on the unsuppressed water reference ^60^, frequency- and phase-correction using probabilistic spectral alignment ^61^, removal of residual water signals using a Hankel singular value decomposition (HSVD) filter ^62^, and modeling of the metabolite peaks ^57,63^. Given that a set of 17 metabolite signals are necessary for the model to accurately fit the spectrum, an appropriate custom basis set was generated using MRSCloud ^64^ with simulated profiles for ascorbate (Asc), aspartate (Asp), creatine (Cr), GABA, glycerophosphocholine (GPC), GSH, glutamine (Gln), glutamate (Glu), myo-I, Lac, NAA, NAAG, phosphocholine (PCh), phosphocreatine (PCr), phosphoryl ethanolamine (PE), scyllo-inositol (sI), and taurine (Tau). The modeled spectral range was from 0.2 to 4.2 ppm and the default Osprey baseline knot spacing of 0.4 ppm was used. Amplitude-ratio soft constraints were applied to the amplitudes of the macromolecules (MM), lipids, and NAAG/NAA peaks as defined in the LCModel manual (http://s-provencher.com/pub/LCModel/manual/manual.pdf). The amplitudes of GABA and the co-edited MM at 3.0 ppm were soft-constrained in a one-to-one ratio, as suggested by previous work ^65^. The water-reference data were modeled in the frequency domain with a dedicated water basis function and a six-parameter model (amplitude, zero- and first-order phase, Gaussian and Lorentzian line broadening, and frequency shift). A summary of the experimental methods and data quality—following the minimum reporting standards in MRS ^65^—is presented in Supplemental Table 1. Although edited MRS methods allow for the detection of numerous metabolites, those with extremely low concentration are less reliable in the context of the study sample size. Therefore, the following metabolites were measured in each participant: NAA, tCho, myo-I, Glu, Gln, and Glx estimated from sum spectra, and GABA, GSH, NAAG, and Lac levels estimated from the difference-edited spectra^48,66^. All metabolite concentrations were estimated after correcting for partial volume effects in each voxel as described in Gasparovic et al. ^67^. Because GABA levels are reported to differ between brain gray and white matter ^68^, GABA levels were alpha-corrected ^69^ and final GABA estimates are reported as GABA+, including the contribution of the modeled co-edited macromolecules^70^. GSH or GABA concentrations equal to 0 were interpreted as evidence of failure to fit and those data points were excluded from further analysis. The normality distribution and homogeneity of variance of metabolites were assessed using the Shapiro-Wilk normality test and F-test, respectively.

## Results

### Participant Characteristics

The AN group was comprised of 13 women with AN-restricting subtype and 3 women with binge/purge subtype. The HC and AN groups were similar in age, ethnicity, and education. The HC group was more racially diverse than the AN group. The AN group weighed less and had a lower BMI relative to the HC group. The AN group had higher EDEQ global score as well as subscales score for eating restraint, eating concern, shape concern, and weight concern scores as well as higher STAI trait anxiety and state anxiety scores relative to the HC group. More women in the AN group relative to the HC group were on psychotropic medications. The mean and standard error of the mean (S.E.M.) or frequency of values of the participant characteristics are displayed in Table 1.

**Table 1.** Characteristics [Mean ± S.E.M. or Frequency (N)] of the women comprising the healthy control (HC) group and anorexia nervosa (AN) group and statistical outcomes of group comparisons.

| Variable | HC Group<br>Frequency (N) or<br>Mean ± SEM | AN Group<br>Frequency (N) or<br>Mean ± SEM | Statistic |
| --- | --- | --- | --- |
| Age (years) | 24.59 ± 2.26 | 24.22 ± 2.89 | $F_{(1,30)}=0.010$, $p=0.919$ |
| Race | Asian (4), Black (1),<br>Multiple (4), Other<br>(2) White (5) | Asian (1), White<br>(14), Multiple (1) | $\chi^2_{(4)}=10.863$, $p=0.028$ |
| Ethnicity | Hispanic (2),<br>Not Hispanic (14) | Hispanic (0),<br>Not Hispanic (16) | $\chi^2_{(1)}=2.133$, $p=0.144$ |
| Education | High School/GED<br>(6),<br>BS/A (6), MS/A (4) | No degree (5),<br>High School/GED<br>(5), AA (1), BS/A<br>(4), MS/A(1) | $\chi^2_{(4)}=8.291$, $p=0.081$ |
| Body Weight<br>(lbs)(Visit 1) | 129.85 ± 4.55 | 104.02 ± 3.99 | $F_{(1,30)}=18.213$, $p<0.001$ |
| BMI (Visit 1) | 21.95 ± 0.44 | 17.58 ± 0.57 | $F_{(1,30)}=37.179$, $p<0.001$ |
| Body Weight<br>(lbs)(Scan Day) | -- | 106.76 ± 3.53 | |
| BMI (Scan Day) | -- | 18.08 ± 0.48 | |
| Duration of Anorexia<br>Nervosa (years) | -- | 6.42 ± 2.86 | |
| EDEQ Global Score | 0.95 ± 0.26 | 3.13 ± 0.43 | $F_{(1,28)}=18.822$, $p<0.001$ |
| EDEQ Eating<br>Restraint Score | 0.83 ± 0.23 | 2.81 ± 0.50 | $F_{(1,28)}=13.010$, $p<0.001$ |
| EDEQ Eating<br>Concern Score | 0.52 ± 0.27 | 2.45 ± 0.40 | $F_{(1,28)}=15.947$, $p<0.001$ |
| EDEQ Shape<br>Concern Score | 1.34 ± 0.34 | 3.87 ± 0.46 | $F_{(1,28)}=19.511$, $p<0.001$ |
| EDEQ Weight<br>Concern Score | 1.09 ± 0.36 | 3.37 ± 0.47 | $F_{(1,28)}=14.756$, $p<0.001$ |
| STAI Trait Score | 39.33 ± 3.03 | 56.93 ± 2.80 | $F_{(1,28)}=18.199$, $p<0.001$ |
| STAI State Score | 38.00 ± 3.13 | 50.00 ± 2.56 | $F_{(1,28)}=8.806$, $p=0.006$ |
| Contraception | Yes (2), No (14) | Yes (2), No (14) | -- |
| Psychotropic<br>Medications | Yes (0), No (16) | Yes (10), No (6) | $\chi^2_{(1)}=14.545$, $p<0.001$ |

SEM: Standard Error of Mean, HC: healthy control, AN: anorexia nervosa, N: sample size, GED: General Education Development, AA: Associate in Arts, BS/BA: Bachelor of Science/Arts, MS/MS: Master of Science/Arts, lbs: pounds, BMI: body mass index, EDEQ: Eating Disorder Examination Questionnaire, STAI: State Trait Anxiety Inventory

### Interoception Measures

On Visit 1 (behavioral assessment of anxiety-to-eat), the AN group reported less hunger, less desire to eat, and more anxiety relative to the HC group. No differences were observed between groups in reported thirst, fullness, and stress. On Visit 2 (MRS assessment of neurometabolites in the dACC), there were no differences in interoceptive measures between groups. The mean and S.E.M. response and statistical outcome for each interoceptive factor is displayed in Supplemental Table 2.

### Anxiety-to-Eat

Main effects of energy density (F_(1,30)_=37.671, p<0.001) and group (F_(1,30)_=27.338, p<0.001) as well as a significant energy density by group interaction (F_(1,30)_=10.602, p<0.003) on anxiety-to-eat ratings were observed. The AN group reported greater anxiety to eat LED foods (p=0.012) and HED foods (p<0.001) relative to responses by the HC group (Figure 2a). Both groups reported greater anxiety to eat HED foods relative to LED foods (HC: p=0.050; AN: p=0.001).

**Figure 2.**
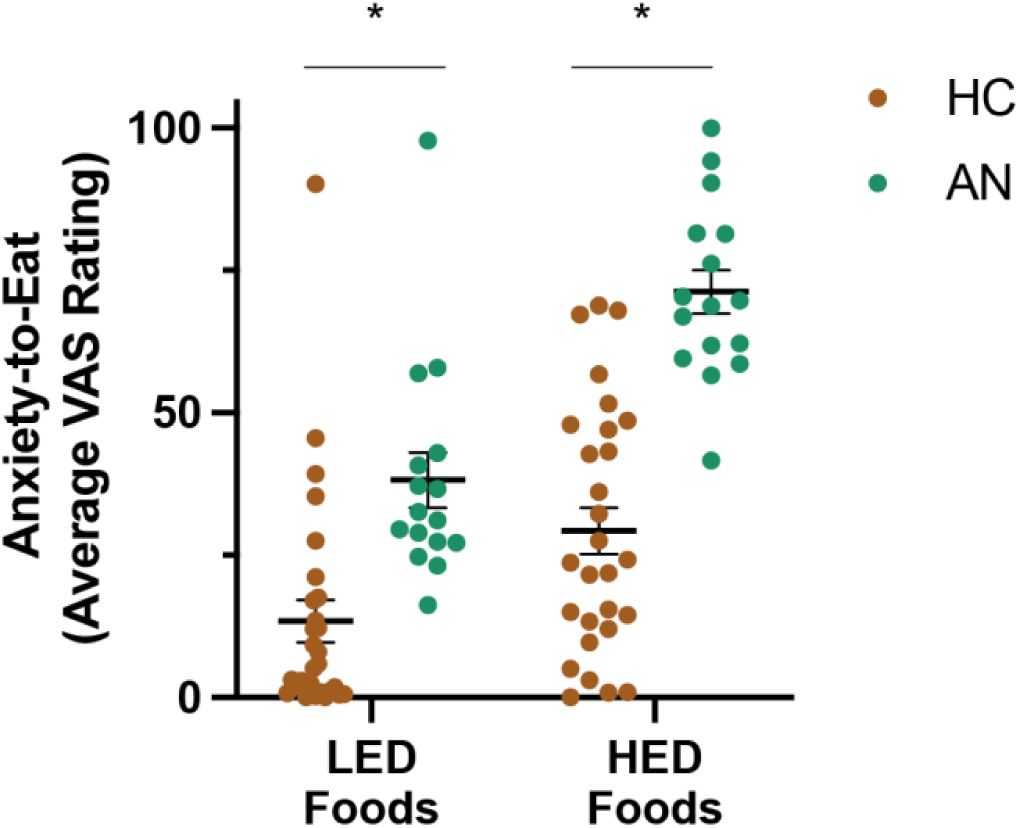
Mean ± S.E.M. anxiety-to-eat ratings for a standard serving size of lower energy dense (LED) and higher energy dense (HED) foods in the healthy control (HC; gold) and anorexia nervosa (AN; cyan) groups. The AN group reported greater anxiety to eat both HED and LED foods relative to the HC group. * represents significant p-value at alpha level of 0.05.

### MRS

MRS data quality was generally high (see Supplemental Table 1). Gray matter (HC: 0.496 ± 0.008, AN: 0.493 ± 0.009; F_(1,30)_=0.088, p=0.768) and white matter (HC: 0.384 ± 0.009, AN: 0.360 ± 0.011; F_(1,30)_=3.068, p=0.090) volume within the voxel were similar between groups, and the AN group had higher CSF levels within the voxel relative to HC group (HC: 0.120 ± 0.003, AN: 0.148 ± 0.010; F_(1,30)_=7.291, p=0.011). The following neurometabolites in the dACC were measured: myo-I, GABA with co-edited macromolecules (GABA+), total choline (tCho), GSH, Glx, NAA, NAAG, Lac, and tCr. Metabolite levels in the dACC region are summarized in Table 2, and their sum spectra overlaid for all participants in each group can be found in Figure 3a. Only myo-I levels differed between groups (Figure 3b, Table 2).

**Figure 3.**
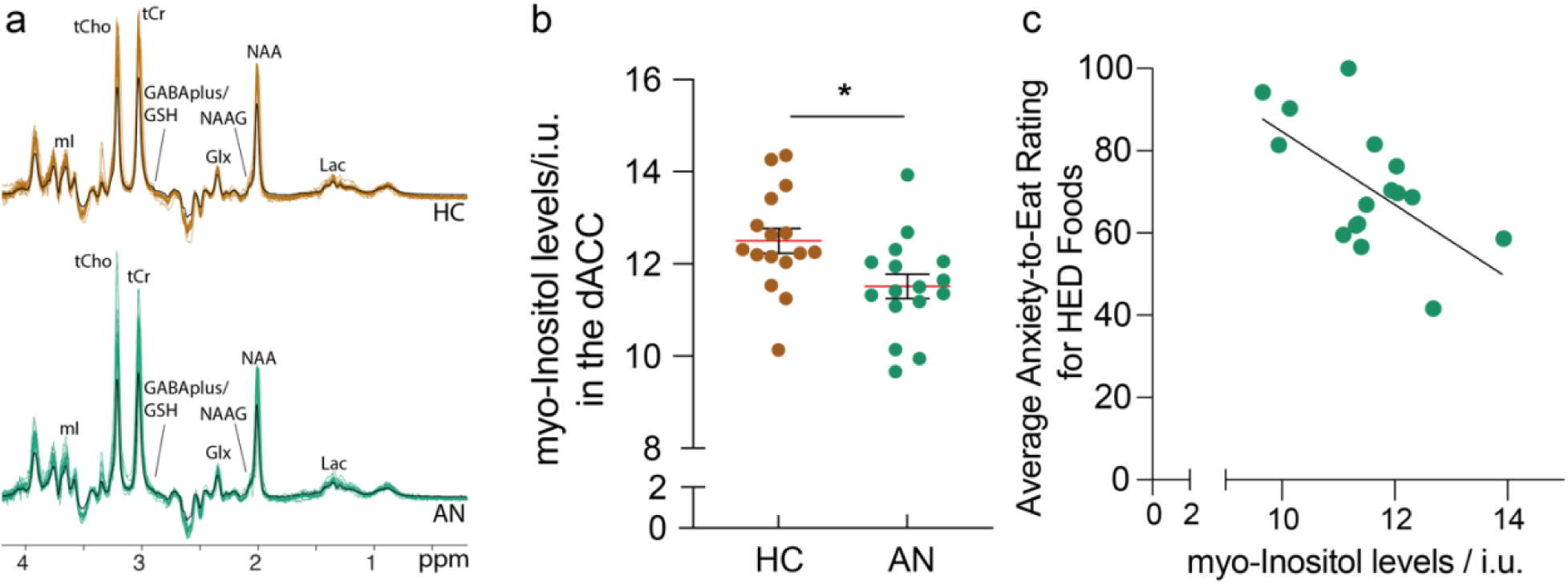
a: In vivo HERCULES data showing the overlaid sum spectra acquired from the dorsal anterior cingulate cortex (dACC) region. The key spectral features of the nine edited metabolites are visible. The black line within each spectrum represents the mean spectrum of each group. The gold spectra are those acquired from the healthy control (HC) group, while the cyan spectra are those acquired from the anorexia nervosa (AN) group. b: Myo-inositol (myo-I) levels in the dorsal anterior cingulate cortex (dACC) as measured by MRS in the healthy control (HC; gold) and anorexia nervosa (AN; cyan) groups. The AN group had lower myo-I levels in the dACC compared to HC. c: Relationship between myo-I levels in the dACC and reported anxiety to eat a standard portion of HED foods in the AN group (cyan). myo-I levels in the dACC negatively predicted anxiety to eat HED foods in the AN group. * regression line represents significant relationship at alpha level of 0.05.

**Table 2.** Metabolites levels in institutional units in the dorsal anterior cingulate cortex (dACC) of women comprising the healthy control (HC) group and the anorexia nervosa (AN) group.

| Neurometabolite | HC Group<br>Mean ± S.E.M. | AN Group<br>Mean ± S.E.M. | Statistic |
| --- | --- | --- | --- |
| myo-I | 12.50 ± 0.27 | 11.51 ± 0.26 | $F_{(1,30)}=6.783$, $p=0.014^*$ |
| GABA+ | 7.74 ± 0.43 | 8.42 ± 0.19 | $F_{(1,30)}=2.097$, $p=0.158$ |
| tCho | 3.10 ± 0.098 | 3.27 ± 0.09 | $F_{(1,30)}=1.770$, $p=0.193$ |
| GSH | 3.63 ± 0.19 | 3.91 ± 0.16 | $F_{(1,30)}=1.203$, $p=0.281$ |
| Glx | 15.59 ± 0.61 | 14.76 ± 0.48 | $F_{(1,30)}=1.128$, $p=0.297$ |
| NAA | 16.17 ± 0.27 | 15.80 ± 0.28 | $F_{(1,30)}=0.921$, $p=0.345$ |
| NAAG | 0.62 ± 0.11 | 0.73 ± 0.12 | $F_{(1,30)}=0.449$, $p=0.508$ |
| Lac | 1.01 ± 0.06 | 1.10 ± 0.08 | $F_{(1,30)}=0.335$, $p=0.567$ |
| tCr | 12.78 ± 0.25 | 12.90 ± 0.19 | $F_{(1,30)}=0.128$, $p=0.723$ |

myo-I: myo-inositol; GABA+: $\gamma$ -aminobutyric acid + co-edited macromolecules; tCho, total Choline; GSH, glutathione; Glx, Glu + Gln; NAA, N-acetyl aspartate; NAAG, N-acetyl aspartyl glutamate; Lac, lactate; tCr, total Creatine; \* represents significant p-value at alpha level of 0.05.

### Relationship Between myo-I Levels in the dACC and Anxiety-to-Eat

myo-I levels in the dACC significantly predicted anxiety to eat HED foods (F_(1,14)_=8.567, p=0.012) in individuals with AN. Specifically, for every unit decrease in myo-I in the dACC, there was an estimated nine-unit increase in anxiety to eat HED foods (β=-9.206, SE(β)=3.145; R^2^=0.397; Figure 3c). General anxiety as reported on the VAS anxiety scale prior to the scan (p=0.528), BMI (p=0.189), duration of illness (p=0.234), and blood glucose levels (p=0.310) were not significant predictors and therefore were not retained in the model. Models could not be fit for anxiety to eat LED foods in AN or anxiety to eat HED or LED foods in HC.

## Discussion

The goal of this project was to characterize anxiety-to-eat, a prominent yet understudied feature of AN psychopathology, and to identify its underlying neurobiological mechanisms. Anxiety-to-eat was higher for HED relative to LED foods in both AN and HC, and anxiety to eat both HED and LED foods was greater in AN relative to HC. Among the nine metabolites examined in the dACC, myo-I levels in the dACC emerged as a significant predictor of anxiety to eat HED foods in individuals with AN but not in HC. This relationship was independent of BMI, duration of illness, blood glucose, and general anxiety state.

These results provide new insight into the clinically challenging feature of eating-related anxiety in AN. Our data complement previous research showing that women with eating disorders report greater anxiety to consume high versus low calorie foods and beverages ^34,71^, and extend these findings to demonstrate that anxiety to eat both HED *and* LED foods is greater in women with AN relative to HC. Moreover, our data demonstrate that HC also report greater anxiety to eat HED relative to LED foods as observed in AN, indicating that anxiety-to-eat is sensitive to energy density and that the *degree* rather than *presence* of anxiety-to-eat is central to AN psychopathology.

Notably, these behavioral data in HC are inconsistent with the report by Ellison et al., ^34^ demonstrating that women without a history of an eating disorder show no such distinction in anxiety to consume high and low-calorie beverages. Differences in methods may underly the inconsistency between the two studies. Ellison et al., ^34^ used images of beverages labeled as high or low calorie and collected anxiety ratings using a Likert scale, whereas we presented images of one standard portion size of 10 HED and 10 LED foods and collected ratings using a VAS that allows for valid across-group comparisons ^72,73^. The variety of the foods presented in our study may have elicited greater variability in ratings of anxiety-to-eat, reflective of differences in food liking, healthiness, or other factors in HC and AN that may not have been captured solely by images of beverages. Alternatively, potential differential factors within the HC group and in sample size between the studies may also account for differences in statistical outcomes; Ellison et al.^34^ assessed six controls and six patients with AN, whereas 16 individuals comprised each group assessed here.

The dACC plays a critical role in threat appraisal and affective decision-making by linking behaviors with positive and negative consequences^74–77^. In AN, HED foods acquire the capacity to elicit negative anticipatory responses including anxiety-to-eat due to fear of body weight or fat gain if consumed. The *perceived* immediate causal consequence of consuming HED foods and resulting heightened emotional state (anxiety-to-eat) may engage the dACC and activate avoidance behavior. This dynamic is not typically observed in individuals without an eating disorder and may explain the absence of a relationship between myo-I levels in the dACC and anxiety-to-eat in HC. If so, myo-I levels in the dACC could serve as a novel biomarker of illness severity independent of BMI and illness chronicity, help to identify individuals vulnerable to AN or eating pathology more generally, and represent a potential therapeutic target. Randomized clinical trials of myo-I supplementation have demonstrated some promising results in other psychiatric conditions with sample sizes ranging from 12-28 individuals ^78–80^.

Our findings replicate prior literature showing lower myo-I levels in the dACC of individuals with AN relative to HC^42,46,47^ and additionally link these lower myo-I levels with the critical phenomenological feature of anxiety-to-eat in AN. Myo-inositol is a carbocyclic sugar and a precursor molecule to numerous phosphatidyl-inositol species in the second-messenger phosphatidyl-inositol systems including adrenergic, serotonergic, and glutamatergic pathways^81,82^ that have been shown to be dysregulated in AN ^83–88^. At least four mechanistic interpretations may help explain lower myo-I in the dACC in AN. First, lower myo-I in the dACC of AN relative to HC may reflect heightened cellular activity or signaling within phosphatidyl-inositol circuits, leading to depletion of the precursor. This would be consistent with prior research showing greater activity in the dACC, or ACC more broadly, to high calorie food images in AN relative to HC^33,34^. Second, the converse may be true; lower levels of myo-I substrate in the dACC of AN relative to HC may reflect lower phosphatidyl-inositol resulting in reduced signaling within such circuits. Third, myo-I is commonly used as a glial marker in the MRS literature. Lower myo-I in the dACC may reflect lower astrocyte density in the PFC, as demonstrated in rodent models of AN ^89,90^. Astrocytes are critical in clearing neurotransmitters from the synapse, and lower numbers of astrocytes may prolong circuit activation due to prolonged neurotransmitter presence at the synapse. Myo-inositol is present in both neurons and glial cells^91^. The inability to distinguish the cell-specificity of measured neurometabolites is a limitation in MRS technology. Fourth, myo-I is primarily derived from de novo synthesis from D-glucose-6-phosphate^92^ and from diet ^93^. Reduced caloric intake or inadequate nutrition for myo-I synthesis may account for lower levels of myo-I in AN relative to HC, although this is less likely given that levels of myo-I were independent of BMI and fasting glucose (in both groups). Similarly, other studies have shown no association between BMI and dACC metrics (i.e., volume, blood flow) during the acute underweight AN state as well as following weight restoration^35,38,43,46,94^.

No differences in the other eight metabolite levels examined in the dACC were observed between groups. This is consistent, although not universally observed, with several studies of similar or larger magnitude investigating concentrations of metabolites in the dACC of underweight individuals with AN^42,43,45,47^ as well as other psychiatric illnesses relative to controls^95–102^. However, an equal number of studies have observed lower levels of NAA^46^, Glx^43,46^, Cr^43^, and Cho^43^ in AN relative to HC. Although the literature is quite limited and the participant characteristics heterogeneous, methodological differences between our study and others including MRI magnet strength^43,44,46^, multivoxel versus single voxel approach^42^, underweight versus weight-restored AN condition^44^, adolescents versus adults^44,46^, and application of our novel editing scheme HERCULES^48^ may account for inconsistencies across the literature. This is the first study to investigate metabolite levels in individuals with AN using HERCULES. HERCULES simultaneously quantifies multiple low-concentration coupled metabolite levels which allows for substantially reduced scan time^66,103^ and treats each target signal independently to enable the extraction of distinct edited spectra from the dataset in the different Hadamard combinations. This provides increased sensitivity and specificity of HERCULES as opposed to PRESS and 2D spectra methods^46,104^

### Limitations

Although comparable to similar observational and MRS studies in the extant AN literature (average 15 participants, range 7-32^105^) and sufficient to identify significant relationships between AN psychopathology and brain chemistry, our sample size was limited. Larger studies are needed to replicate and extend these findings. The MRS method used here corrected for water and metabolite relaxation constants in tissues based on the assumption that T1/T2 values are uniform as indicated in Wansapura et al.,^106^. Changes in water and metabolite relaxation rates have been observed with age ^55,107,108^ and served as the justification for age-matching the groups. Both amenorrhea and use of psychotropic medications are common in AN, but their impact on MRS metrics remain unclear. While previous studies have found no effect of the menstrual cycle on myo-I levels^109^ or medication effects on MRS metrics in individuals with psychiatric conditions (e.g., OCD, TD)^97^, their effects cannot be entirely dismissed. For both ethical and practical reasons, participants did not discontinue these medications before participation in this study. Notably, we found no differences in myo-I levels in women with AN receiving psychotropic medications versus those not currently receiving such treatment.

### Conclusion

The degree of anxiety-to-eat differs between women with AN compared to HC and was negatively correlated with myo-I in the dACC in women with AN but not in HC in this study. Although unrelated to BMI, duration of illness, or general anxiety, myo-I levels in the dACC were negatively associated with anxiety-to-eat, raising the question of whether low myo-I in the dACC in individuals with AN mechanistically underlies anxiety-to-eat. Future longitudinal studies in humans and work in preclinical models are needed to replicate and extend these findings, to clarify whether the relationship between myo-I and anxiety-to-eat changes with weight restoration and to determine if anxiety-to-eat is modified by dietary supplementation with myo-I or is predictive of treatment response.

## Author Contributions

Author Contributions: Conceptualization (KRS, RAEE); Data collection and curation (KRS, SG, YS); Formal analysis (CWD, KRS, YS); Funding acquisition (KRS); Writing - original draft (KRS, YS); Writing - review & editing (AG, CWD, KRS, RAEE, SG, YS). All authors have approved of the final article.

## Acknowledgements

The authors would like to thank the research participants and MR technicians Terri Brawner, Kathleen Kahl, and Ivana Kusevic for assistance with MR data collection. A preprint of the manuscript was posted on bioRxiv.

## Funding

Funding for this project was supported by the National Institute of Mental Health K01MH127178, Dalio Philanthropies, R21HD100869, R01EB023963, R01EB032788, and R01EB016089.

## Competing Interests

Dr. Guarda reports receiving royalties from UpToDate. All other authors report no biomedical financial interests or potential conflicts of interest.

## Notes

### Competing Interest Statement

The authors have declared no competing interest.

### Summary of Updates

Figures are revised. Discussion are revised.

