## Supplemental Table 1; Supplemental Table 2 for "Myo-inositol in the Dorsal Anterior Cingulate Cortex is Associated with Anxiety-to-Eat in Anorexia Nervosa"

**Supplemental Material**

Supplemental Table 1: Summary following minimum reporting standards in MRS generated in Osprey See Lin et al. 'Minimum Reporting Standards for in vivo Magnetic Resonance Spectroscopy (MRSinMRS): Experts' consensus recommendations. NMR in Biomedicine. 2021;e4484. [doi.org/10.1002/nbm.4448](https://doi.org/10.1002/nbm.4448)

| 1.Hardware | |
| --- | --- |
| a. Field strength [T] | 3 T |
| b. Manufacturer | Philips |
| c. Model (software version if available) | R 5.7.1 |
| d. RF coils: nuclei (transmit/receive), number of channels, type, body part | ^1^H, 32 channel, head |
| e. Additional hardware | - |
| 2. Acquisition | |
| a. Pulse sequence | HERCULES (Johns Hopkins University Patch) |
| b. Volume of interest (VOI) locations | dACC |
| c. Nominal VOI size [mm^3^] | dACC:30 x 30 x 30 mm^3^ |
| d. Repetition time (TR), echo time (TE)[ms] | TR 2000 ms, TE 80 ms |
| e. Total number of averages per spectrum  i. Number of averaged spectra per subspectrum | 320 total averages with 80 averages per subspectrum |
| f. Additional sequence parameters  i. Editing pulse parameters | F1: 2000 Hz, 2048 points  GABA at 1.9 ppm, GSH at 4.56 ppm |
| g. Water suppression method | VAPOR |
| h. Shimming method, reference peak, and threshold of acceptance of shim chosen | 1^st^ and 2^nd^ shimming pencil beam, water |
| i. Trigger or motion correction | No trigger or active motion correction |
| 3. Data analysis methods and outputs | |
| a. Analysis software | Osprey 2.4.0 |
| b. Processing steps deviating from Osprey | Final alignment of the averaged sub-spectra by  minimizing the choline (not water) peak; co-edited MMs at 3 ppm were modelled using the  “1to1GABAsoft” model |
| c. Output measure | tCr, rawWaterScaled, CSFWaterScaled, TissCorrWaterScaled |
| d. Quantification references and assumptions, fitting model assumptions | Asc, Asp, Cr, GABA, GPC, GSH, Gln, Glu, mI, Lac, NAA, NAAG, PCh, PCr, PE, scyllo-inositol, Tau, 8 MM basis functions in the sum spectrum (MM_0.94_, MM_1.22_, MM_1.43_, MM_1.70_, MM_2.05_, Lip09, Lip13, Lip20) Fitting method: Osprey baseline knot spacing 0.4 ppm |
| 4. Data quality | |
| HERCULES, dACC | |
| a. SNR (Cr), linewidth (Cr) [Hz, OFF spectra] | SNR 233 ± 40, linewidth 5.05 ± 0.66 Hz |
| c. Quality measures of postprocessing model fitting (Mean Relative Amplitude Residual (Residual/Noise))  sum  diff1  diff2 | 22.35 ± 22.18  5.20 ± 1.42  5.35 ± 1.76 |
| d. Mean spectrum created with OspreyOverview | Figure 1 |

T: Tesla, RF: radiofrequency, HERCULES: Hadamard Editing Resolves Chemicals Using Linear-combination Estimation of Spectra, mm: millimeter, ms: millisecond, Hz: frequency, ppm: parts per million, VAPOR: Variable Power radiofrequency pulses with Optimized Relaxation delays, dACC: dorsal anterior cingulate cortex, SNR: signal-to-noise ratio, GABA: γ-aminobutyric acid, GSH: glutathione, Asc: ascorbate, Asp: aspartate, Cr: Creatine, GPC: glycerophosphocholine, Gln: glutamine, Glu: glutamate, mI: myo-inositol, Lac: lactate, NAA: N-acetyl aspartate, NAAG: N-acetyl aspartyl glutamate, PCh: phosphocholine, PCr: phosphorcreatine, PE: phosphoryl ethanolamine, Tau: taurine, MM: macromolecules, Lip: lipid.

Supplemental Table 2. Mean ± S.E.M. response for interoceptive factors in women comprising the healthy control (HC) group and anorexia nervosa (AN) group.

| **Variable** | **HC Group** | | **AN Group** | | **Statistic** | |
| --- | --- | --- | --- | --- | --- | --- |
| Visit 1 (Behavioral Assessment of Anxiety-to-Eat) | | | | | | |
| Hunger | 48.37 ± 4.26 | | 31.75 ± 6.06 | | F_(1,30)_=5.033, p=0.032* | |
| Thirst | 49.75 ± 5.23 | | 51.88 ± 5.07 | | F_(1,30)_=1.169, p=0.288 | |
| Fullness | 37.50 ± 3.56 | | 48.81 ± 6.72 | | F_(1,30)_=2.214, p=0.147 | |
| Desire to Eat | 55.69 ± 5.69 | | 32.00 ± 5.63 | | F_(1,30)_=8.749, p=0.006* | |
| Anxiety | 25.88 ± 6.44 | | 49.94 ± 5.91 | | F_(1,30)_=7.589, p=0.010* | |
| Stress | 31.44 ± 7.84 | | 49.44 ± 5.77 | | F_(1,30)_=3.420, p=0.074 | |
| Visit 2 (MRS Scanning Protocol) | | | | | | |
| Hunger | | 50.13 ± 7.23 | | 36.63 ± 5.96 | | F_(1,30)_=2.078, p=0.160 |
| Thirst | | 49.56 ± 6.29 | | 45.44 ± 6.40 | | F_(1,30)_=0.212, p=0.649 |
| Fullness | | 28.50 ± 5.11 | | 41.63 ± 7.21 | | F_(1,30)_=2.205, p=0.148 |
| Desire to Eat | | 50.57 ± 7.20 | | 38.06 ± 5.76 | | F_(1,30)_=1.838, p=0.185 |
| Anxiety | | 28.50 ± 6.60 | | 41.31 ± 6.44 | | F_(1,30)_=1.931, p=0.175 |
| Stress | | 29.19 ± 7.44 | | 41.38 ± 5.52 | | F_(1,30)_=1.731, p=0.198 |

SEM=Standard Error of Mean; HC=healthy control; AN=anorexia nervosa; * represents significant p-value at alpha level of 0.05.
